# MPTHub: an open-source software for characterizing the transport of particles in biorelevant media

**DOI:** 10.1101/2021.09.15.460434

**Authors:** Leandro Gabriel, Helena Almeida, Marta Avelar, Bruno Sarmento, José das Neves

**Author notes:** These authors contributed equally to this work (Leandro Gabriel, Helena Almeida).

## Abstract

The study of the transport of particles in different environments plays an essential role in understanding interactions with humans and other living organisms. Importantly, obtained data can be directly used for multiple applications in fields such as fundamental biology, toxicology or medicine. Particle movement in biorelevant media can be readily monitored using microscopy and converted into time-resolved trajectories using freely available tracking software. However, translation into tangible and meaningful parameters is time-consuming and not always intuitive. Thus, we developed a new software – MPTHub – as an open-access, stand-alone, user-friendly tool for the rapid and reliable analysis of particle trajectories extracted from video microscopy. The software was programmed using Python and allowed to import and analyze trajectory data, and export relevant data such as individual and ensemble time-averaged mean square displacements and effective diffusivity, and anomalous transport exponent. Data processing was reliable, fast (total processing time of less than 10 sec) and required minimal memory resources (up to a maximum of around 150 MB in RAM). Demonstration of software applicability was conducted by studying the transport of different polystyrene nanoparticles (100-200 nm) in mucus surrogates. Overall, MPTHub represents a freely available software tool that can be used even by unexperienced users for studying the transport of particles in biorelevant media.

## 1. Introduction

Nanotechnology and materials science have come a long way over the last few decades, contributing not only to major scientific breakthroughs, but also to tangible everyday solutions that are flooding the global market [1]. As a consequence, humans are increasingly exposed to nanomaterials, either intentionally (*e.g*., in case of nanomedicine use) or inadvertently (*e.g*., resulting from occupational contact with nanopollutants). Outcomes of such events are dependent on complex interactions established at key gateways of the human body, namely mucosal sites, which need to be thoroughly recognized and studied [2]. Transport across different natural barriers and media are of particular importance as it largely determines the fate of nanomaterials. Indeed, the possibility to study motion of molecules and supramolecular structures at the nano- and micro-scales opened new doors to the understanding of the dynamics and interactions of such entities in complex biological environments [3-6]. Methods for particle tracking generally fit into one of the following two categories: (i) methods that measure the ensemble average dynamics of particles – such as fluorescence recovery after photobleaching (FRAP), fluorescence correlation spectroscopy (FCS) or dynamic light scattering (DLS) –, and (ii) methods that are able to discriminate single particle movement. Despite enhanced signal due to bulk emission combined with high temporal resolution provided by ensemble methods, these are known for overlooking important information regarding heterogeneous particle populations or media [7]. Multiple particle tracking (MPT) is a well-established technique that enables tracing individual molecules, particulates or even microorganisms at a high spatiotemporal resolution in complex biological systems such as the intracellular environment, extracellular matrix or biological fluids (*e.g*., blood or mucus) [8]. This powerful technique has been widely used in the biomedical field in order to assess the intracellular transport and kinetics of proteins, lipids and molecular motors [9], study cell migration and mobility of pathogens [10, 11], determine the mechanical properties of isolated nuclei in living cells [12], or characterize the microstructure and rheological profile of mucus and other viscoelastic fluids [13-15], to name only a few applications. In the particular case of nanomedicine, MPT allowed understanding the transport of nanocarriers in biological media and helped devising engineering strategies that could be useful in overcoming natural barriers to drug delivery and, thus, enhance therapeutic outcomes [16-18].

Details on principles, procedures and even equipment for conducting MPT experiments are quite simple and readily accessible [8]. The processing cascade of MPT includes the acquisition of real-time videos of typically fluorescently labeled molecules or structures in biorelevant media using a microscope and a high sensitivity camera. Different processing algorithms available as plugins in common image processing programs can be employed to analyze videos and identify particles, plot their frame-by-frame coordinates and define individual trajectories [19]. For example, the MosaicSuite package [20], NanoTrackJ [21] and TrackMate [22] for ImageJ/Fiji are popular and free to use plugins. Quantitative raw data in the form of biologically relevant parameters can then be extracted from obtained trajectories, either to understand the diffusive behavior of particles or characterize the 3D network and rheological properties of the specimen in which these are embedded. While the ability to analyze motion at the single particle level provides a valuable body of information that is not possible to attain with purely ensemble techniques, manual handling of generated large datasets constitute a prone to error and time-consuming process [23]. Various available software have been shown useful in aiding this task [24-27]. However, these solutions offer limited possibilities for data processing customization, need advanced computing skill, or require payment of a license (self or that of an ancillary software such as MATLAB). Software provided with analytical equipment can also undertake MPT analysis (e.g., NIKON NIS Elements or Malvern NTA software), although their use is limited to specific hardware [28, 29]. Similar generic open-source instrumentation software is available, but the number of compatible equipment, namely microscopes, is still limited [30].

This work details on the development of an open-access, stand-alone, user-friendly software named MPTHub for the rapid and reliable analysis of particle trajectories extracted from video microscopy. The source code and latest version of the software are available for download at https://github.com/lassisg/mpthub. The current version of MPTHub (v. 1.0.1) is available under Git version control as an open-source software for Microsoft Windows (GNU General Public License v. 3.0). The relevance of the software was further demonstrated by characterizing the transport of 100-200 nm fluorescent polystyrene nanoparticles (NPs) in intestinal mucus surrogates.

## 2. Materials and methods

### 2.1. Materials

Red fluorescent carboxylate-modified polystyrene (COOH-PS) NPs with nominal mean diameter of 100 nm or 200 nm were purchased from Molecular Probes (Eugene, OR, USA). Phosphate buffered saline (PBS) tablets and purified type II mucin from porcine stomach were purchased from Sigma (St. Louis, MO, USA), and 1.5 × 1.6 cm gene frames (65 µL) were from Fisher Scientific (Hampton, NH, USA), and Amicon Ultra-0.5 mL (100 kDa MWCO) centrifugal filters were from Merck Millipore (Burlington, MA, USA). Poloxamer 407 – a poly(ethylene glycol)-poly(propylene glycol)-poly(ethylene glycol) (PEG-PPG-PEG; average MW of 9.8 to 14.6 kDa) – was a kind offer from BASF (Ludwigshafen, Germany). All other reagents and materials were of analytical grade or equivalent.

### 2.2. Software development, data processing and performance

MPTHub was developed using Python (v. 3.9.5, available at https://www.python.org/) to enable researchers to easily and rapidly perform particle tracking analysis without compromising the reliability of generated results, as often associated with manual handling of large scientific datasets [31]. Other packages were further used for the development of MPTHub, namely PySide6 for Graphical User Interface (GUI; v. 6.1.1, available at https://pypi.org/project/PySide6/), Pandas for data manipulation (v. 1.2.4, available at https://pandas.pydata.org/), Numpy for mathematical operations (v. 1.20.3, available at https://numpy.org/), MatplotLib for plotting (v. 3.4.2, available at https://matplotlib.org/), XlsxWriter for spreadsheet generation (v. 1.4.3, available at https://pypi.org/project/XlsxWriter/) and TrackPy for particle tracking analysis [32] (v. 0.5.0, available at https://soft-matter.github.io/trackpy/v0.5.0/).

The software was designed in order to comply with three major sequential tasks: (i) import data extracted from microscopy video files, (ii) analyze data, and (iii) export results. The possibility to adjust configurations and proceed with re-analysis was also considered. Input files (.csv format) were generated from microscopy videos, and included data for two-dimensional Cartesian coordinates per frame (see sub-section 2.3.3 for details on file acquisition and pre-processing). MPTHub requires minimal initial configuration settings by the user (*e.g*., metric to digital units conversion), and provides several default values that can be changed. These include particle size, trajectory length cutoff (minimum number of consecutive frames that define valid trajectories), total length of the video in frames, pixel and temporal resolution, lag time or time scale (*τ*) for analysis, and sample temperature during video acquisition. The software was developed in order to allow users to interact and define performed actions performed at multiple points of data processing (**Figure 1**).

**Figure 1.**
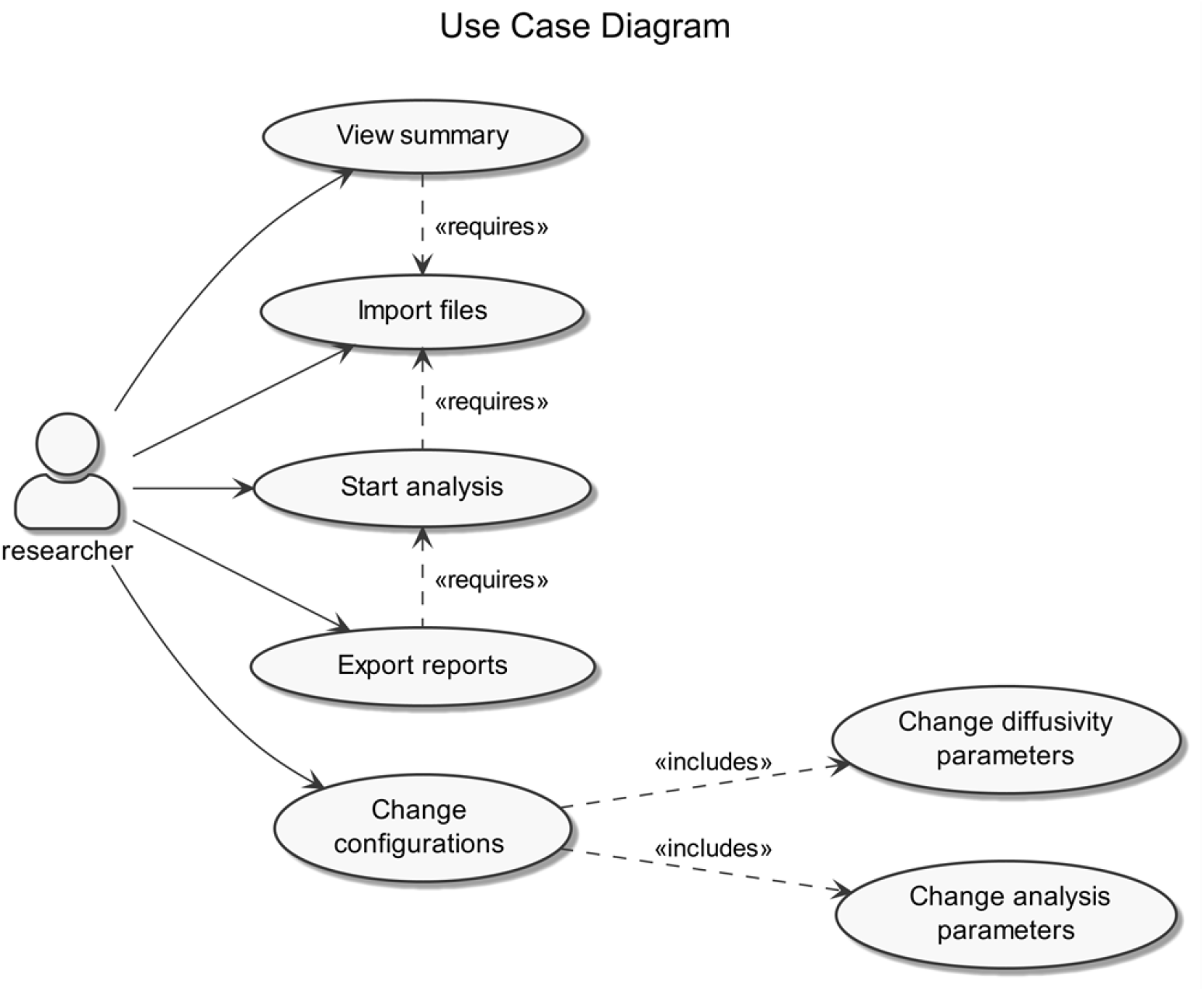
MPTHub Use Case diagram.

Upon analysis, the software was designed to allow calculating the following parameters: (i) time-averaged mean square displacements (MSD or ⟨Δ*r*^2^(*τ*)⟩), (ii) time-dependent diffusion coefficient or effective diffusivity (*D*_eff_), (iii) *D*_w_/<*D*_eff_> ratio, and (iv) anomalous transport exponent (*α*). MSD values for individual trajectories were determined using the following equation:

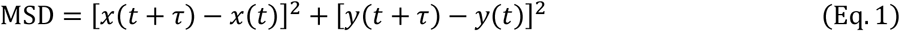

where *x* and *y* denote coordinates between consecutive *τ* intervals [33]. For a particle diffusing in a simple viscous liquid, the MSD was related to *D*_eff_ according to:

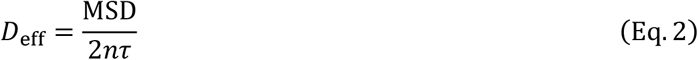

where *n* refers to dimensionality (*n*=2 in a two-dimensional system) [33]. Ensemble averages of MSD (<MSD>) and *D*_eff_ (<*D*_eff_ >) were further calculated and plotted against lag time. The ⟨*D*_eff_⟩ at a given lag time can be compared with the theoretical diffusion coefficient of spherical particles in water (*D*_w_), as calculated by the Stokes-Einstein equation:

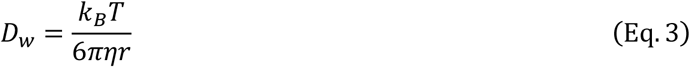

where *k*_*B*_ is the Boltzmann’s constant, *T* the temperature, *η* the fluid viscosity, and *r* the particle hydrodynamic radius. The ratio between *D*_*w*_ and *D*_eff_ provides a relative measure of transport impairment of particles in a specific medium as compared to water. In many cases the diffusion of particles follows a power law scaling that allows determining α according to:

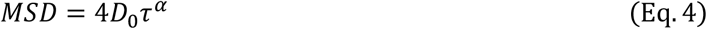

where *D*_0_ is the time-independent diffusion coefficient. Additionally, the value of *α* can be used to classify particles as immobile (*α* < 0.2), sub-diffusive (0.2 ≤ *α* < 0.9), diffusive (0.9 ≤ *α* < 1.2), or active (*α* ≥ 1.2) [34, 35]. The classification as diffusive corresponds to Brownian motion as predicted by the Stokes-Einstein relation [33].

The performance of MPTHub was also assessed using different sets of input files (two or four files) featuring a variable number of valid trajectories (100, 500 or 1,100). Evaluated performance parameters included computing time and memory usage (defined as RAM peak value achieved during processing) for a full cycle of analysis, which included data import, data analysis and results export. Testing was performed in triplicate by using different hardware for replicates (details of each machine are presented in *Supplementary Information, Table S1*).

### 2.3. Application of MPTHub

#### Processing of polystyrene nanoparticles

Fluorescent COOH-PS NPs were washed once with ultrapure water using centrifugal filters (4,300 rpm, 10 min) in order to remove traces of sodium azide. NPs were then resuspended in the same solvent at concentrations varying between 5 and 20 mg.mL^-1^, and dispersions were stored at 4 °C until further use. Coating with poloxamer 407 was achieved by incubating 100 nm NPs in a 1% (*w/v*) polymer solution overnight. NPs were then washed trice before usage, as detailed above. The hydrodynamic diameter, polydispersity index (PdI) and zeta potential were determined by dynamic light scattering and laser Doppler electrophoresis using a Zetasizer Nano ZS (Malvern Instruments, Malvern, UK). NPs were diluted between 0.1 and 0.01 mg.mL^-1^ in 10 mM sodium chloride (pH 5.8) and measurements were carried out at 25 °C.

#### Preparation of mucus surrogates

An intestinal mucus surrogate was prepared by modifying a previously described recipe [34]: mucin was reconstituted in isotonic PBS (pH 7.4) at 30 mg.mL^-1^ or 50 mg.mL^-1^ with the aid of a vortex mixer. The mucus simulant was left to equilibrate for 30 min at room temperature before 65 µL being gently transferred into a frame chamber mounted on a glass microscope slide. Two microliters of NPs dispersed in PBS were added on top of the mucus surrogate and left to equilibrate for 1 h at room temperature after sealing the chamber with a coverslip. The final concentration of NPs in the surrogate was 9.0×10^−6^% (*w/v*) or 7.5×10^−5^% (*w/v*) for 100 nm and 200 nm NPs, respectively.

#### Microscope configuration and video acquisition

The transport of fluorescent NPs was assessed by analyzing their trajectories in either mucus surrogates or water. MPT videos were recorded by using a Hamamatsu ORCA-Flash4.0 digital CMOS camera (Hamamatsu, Japan) mounted on a Leica DMI6000 inverted epifluorescence microscope (Wetzlar, Germany) equipped with a 63×/1.30 NA GLYC-immersion objective. Videos (512 × 512 pixels, 16 bit, 20 sec duration) were collected with the LAS X software at a temporal and pixel resolution of 33 milliseconds and 0.159 µm, respectively. Tracking resolution of approximately 20 nm was determined by calculating displacements of NPs immobilized in a strong adhesive. Acquired video files (.lif format) were pre-processed through background subtraction using ImageJ/Fiji (v. 2.1.0, available at https://imagej.net/software/fiji/), and two-dimensional Cartesian coordinates of particle centroids were detected at sub-pixel resolution using the MosaicSuite 2D/3D single-particle tracking plugin developed by Sbalzarini and Koumoutsakos [20]. Data were exported as .csv files and analyzed with the MPTHub software. Trajectories with at least 100 consecutive frames were considered valid for transport analysis. Three independent experiments, each collecting the trajectories of a minimum of one hundred NPs, were conducted for different conditions.

## 3. Results and discussion

### 3.1. MPTHub development, features and performance

#### Software programming and workflow

Python was selected as programming language due to its open access and open source status, reliability/stability, relatively fast running time and low memory consumption, reduced development time, and condensed amount of generated code as compared to competing software [36]. MPTHub was designed in order to perform three sequential main tasks that can be regarded as independent operational modules with particular processing sets: (i) data import, (ii) data analysis and (iii) results export. First, data importing starts upon the user’s command for loading one or multiple input files (.csv format), which were previously generated by the MosaicSuite ImageJ plugin. Content and format compatibility is then checked upon selection of the ‘import’ command, and data are sanitized for compliance with TrackPy requirements for content organization. The software generates and displays a summary of total and valid trajectories per input file. Upon user’s command, valid trajectories are processed using TrackPy for calculating relevant transport parameters. Exporting command generates and saves external reports that can be readily accessed by the user.

#### Graphical user interface and utilization

The GUI of MPTHub features a simple, user-friendly and flexible design, allowing intuitive use even from unexperienced individuals (**Figure 2**). The launch screen is divided from top to bottom into command menus, fast-access toolbars, and imported file listing and summary of total and valid trajectories. The software allows simultaneous importing and pre-processing of multiple input files (.csv format) in order to provide a summary of total and valid trajectories. The software allows customizing configurations regarding analysis and transport mode diffusivity ranges according to specific needs (**Figure 3**). The user can then select which files should be analyzed, including for iterations. Three data reports are generated as .xlsx format spreadsheets (named as ‘Individual Particle Analysis’, ‘Transport Mode Characterization’ and ‘Stokes-Einstein Calculations’) in order to allow flexible handling of generated content according to the requirements of individual users (*Supplementary Information, Figures S1* to *S9*). Data resulting from software analysis are only temporarily stored in local hardware, unless exported by the user to a folder of choice. The last initial user input configurations are saved for convenience as a small SQLite database file (v. 3.36.0, available at https://www.sqlite.org/) that is created upon the first run of MPTHub. This embedded system does not require installation and is characterized by its portability [37].

**Figure 2.**
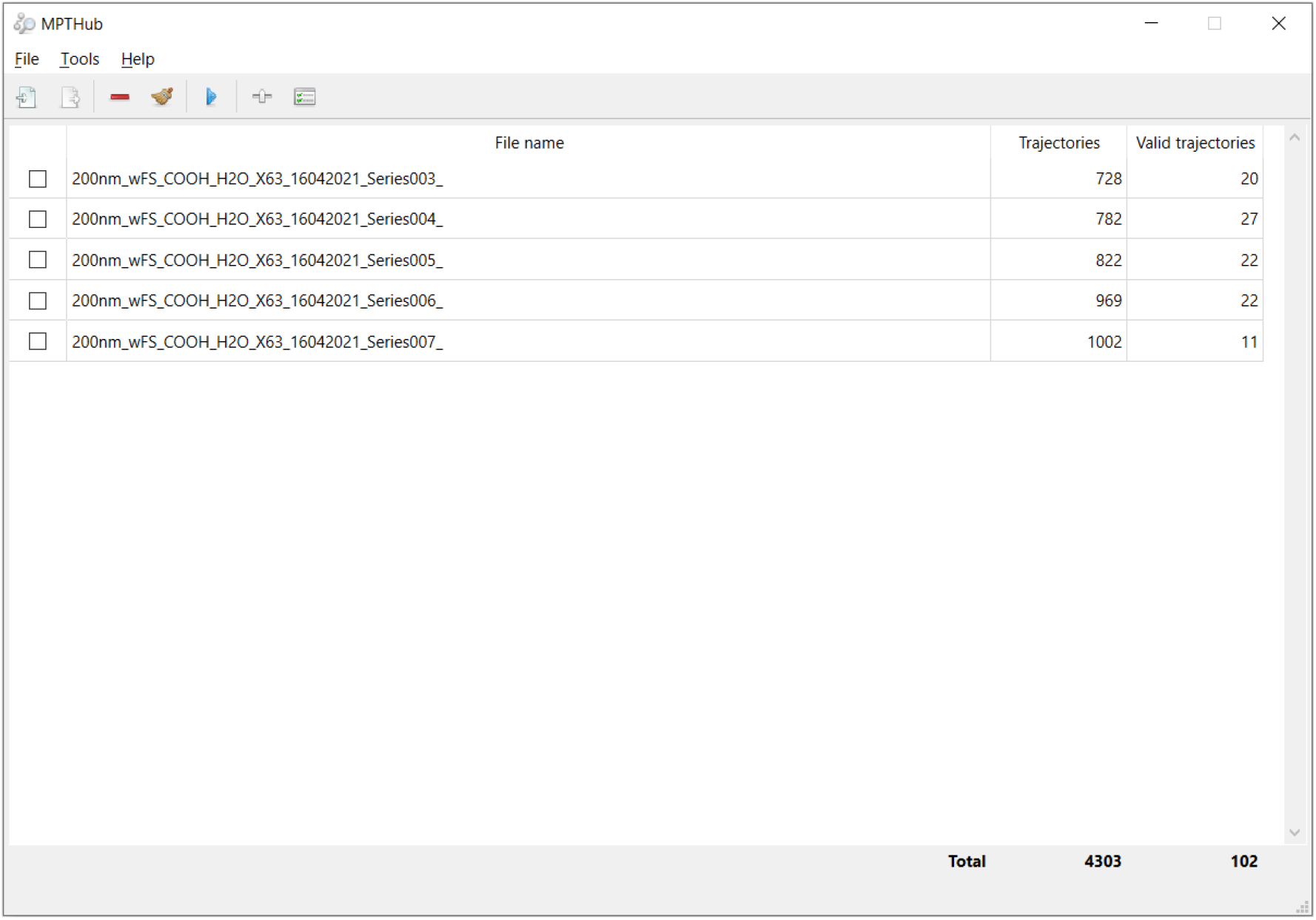
Launch screen capture of MPTHub. Multiple input files are presented (file names, total trajectories and valid trajectories according to defined settings). Included icons are used under CC BY 3.0 license (Copyright 2021, Yusuke Kamiyamane, available at https://p.yusukekamiyamane.com/).

**Figure 3.**
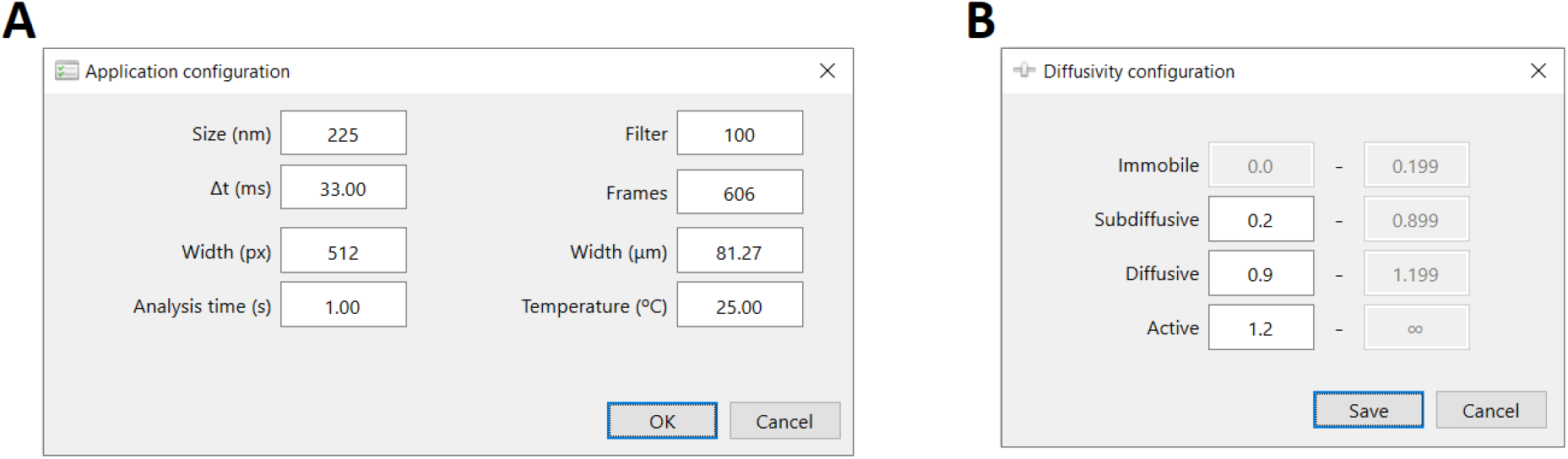
Input configuration parameters of MPTHub. Panels used by the user for defining **(A)** experimental conditions (particle size, temperature), input video properties (field width in pixels and micrometers, temporal resolution, and total length in frames), and analysis parameters (minimum number of consecutive frames needed to define valid trajectories (filter), and analysis time); and **(B)** range for the anomalous diffusion exponent in order to define transport mode.

#### Software performance

One of the main purposes for developing MPTHub was to abbreviate the tedious and time-consuming process involved in manual data curation and processing. This usually involves the creation and use of standard template spreadsheet files, taking hours to days depending on the number of files requiring analysis and user’s experience [34, 35]. Generically, MPTHub reduced the total computing time for a full cycle of data processing for multiple files (typically up to 20), including importing and exporting, to less than 10 sec. In order to provide a more objective review of computing performance MPTHub and to better understand the impact of data volume on its performance, we estimated the computing time and memory usage required for processing two sets of input files at different levels of valid trajectories (adjusted by defining suitable analysis parameters). The use of different machines (*Supplementary Information, Table S1*) with various processing capabilities had minimal impact in terms of RAM peak usage, with relative standard deviation (RSD) values varying between 0.5% and 2.7% for all considered input files/valid trajectories. These values were higher for computing time values of a full cycle of data processing (RSD between 45.5% and 61.2%). Still, in practical terms, such differences are likely irrelevant and do not limit the use of MPTHub even in less resourceful standard computers. Total processing time of MPTHub increased with higher amounts of input data (**Figure 4A**). Still, maximum total processing time was only 8.6 sec under unusually high input data packages (four files/1,100 valid trajectories). A similar trend was noted for data analysis and results export stages. However, import time was mostly affected by the number of input files, not valid trajectories, since processing at this stage is proportional to in-memory data storage. RAM usage was relatively constant along different processing stages, and was to be mostly dependent on the number of files being processed rather than the amount of valid trajectories (**Figure 4B**). The utilization of RAM peaked at roughly 100 MB and 150 MB when processing two or four files, respectively, from a value of around 87 MB at standby. This marginal increase and relative stable use of memory resources throughout import, analysis and export stages indicate that the choice for a software design that does not store data locally had little impact in RAM usage.

**Figure 4.**
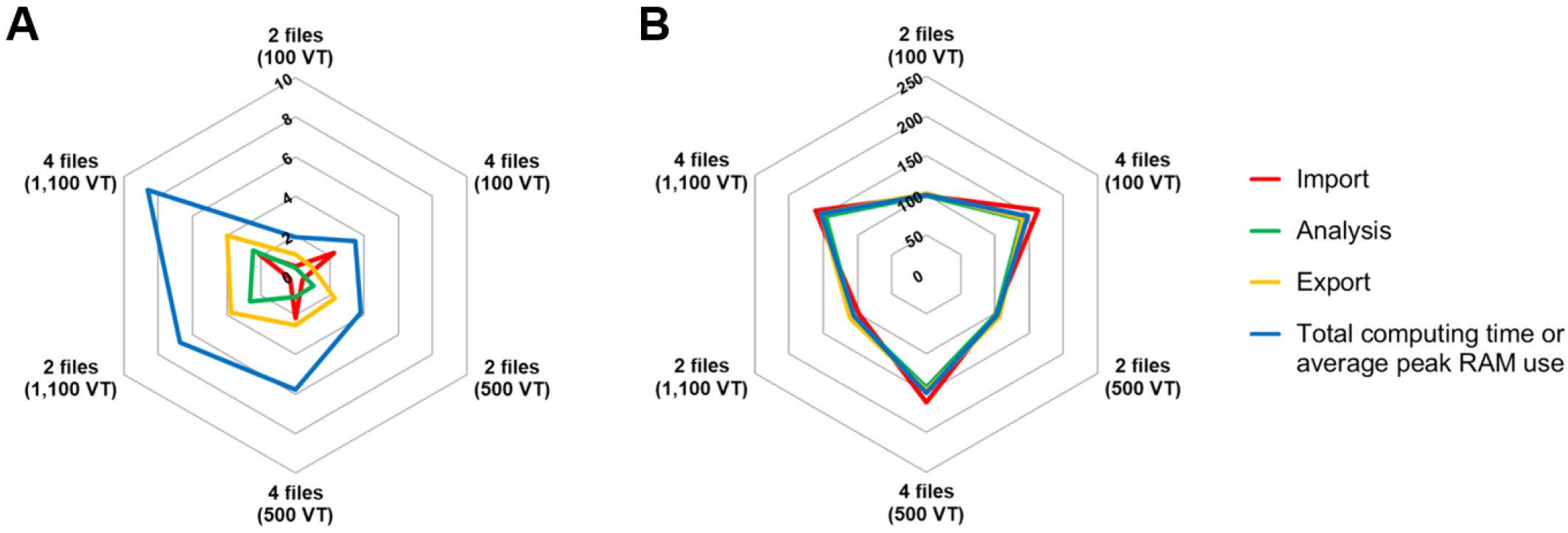
Performance analysis of MPTHub. **(A)** Computing time (in sec), and **(B)** RAM peak use (in MB) when processing two or four input data files containing 100, 500 or 1,100 valid trajectories (VT). Results are presented as mean values (*n* = 3).

### 3.2. Application of MPTHub

Demonstration of the potential of MPTHub was conducted by analyzing the motion of fluorescent COOH-PS NPs in intestinal mucus surrogates. These particles were selected due to the substantial quanta emitted upon excitation, which enables the reduction of exposure time and therefore potential dynamic errors. NPs were monodisperse and presented hydrodynamic values close to the nominal diameter provided by the supplier (**Table 1**). Changes in diameter for 100 nm NPs modified with poloxamer were negligible, while the increase in zeta potential to near neutral values suggests extensive non-covalent coating with the triblock copolymer. The formation of a dense, low MW PEG corona is known to confer mucus-penetrating properties to similarly sized NPs [38].

**Table 1.**
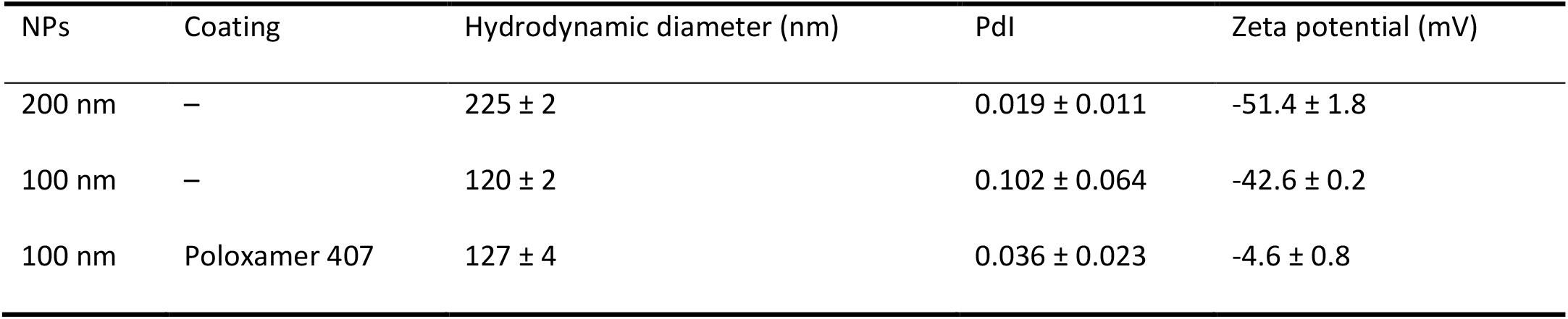
Properties of COOH-PS NPs. Results are presented as mean ± SD (*n* = 3).

The transport properties of NPs in different media were analyzed using the proposed software. We started by comparing the motion of 200 nm COOH-PS NPs in water and mucus surrogate containing 3% mucin (**Figure 5A**; and *Supplementary Video 1* and *Supplementary Video 2*). Behavior of NPs in water was random (typical of Brownian motion) and relatively constant regarding spanned length. Conversely, the transport of NPs in surrogate mucus was quite variable, ranging from highly restricted to nearly unhindered as compared to water. This seems to denote a highly heterogeneous structure of the simulated medium, which is consistent with the properties of native mucus [39]. Differences in NP transport of 200 nm COOH-PS NPs were further confirmed and quantified by ensemble (**Figure 5B-C**) and individual (*Supplementary Information, Figures S10* and *S11*) analysis of MSD and *D*_eff_. Values of <MSD> increased gradually with time scale, while those for <*D*_eff_> remained constant, thus suggesting that particles experienced thermally driven Brownian motion [33]. Also, <MSD> for 200 nm COOH-PS particles for *τ* = 1 sec was almost 7-fold lower in mucus surrogate than for equivalent particles in water. For the same time scale, transport in simulated mucus was nearly 6-fold lower (*D*_w_/*D*_0_ = 5.8) than what would be expected in water, according to the Stokes-Einstein equation. Transport in water indicates an overall good agreement with theoretical diffusivity (*D*_w_/*D*_0_ = 0.9), with the slight deviation likely resulting from MSD variation at longer times scales (considering the length of the trajectories) [40].

**Figure 5.**
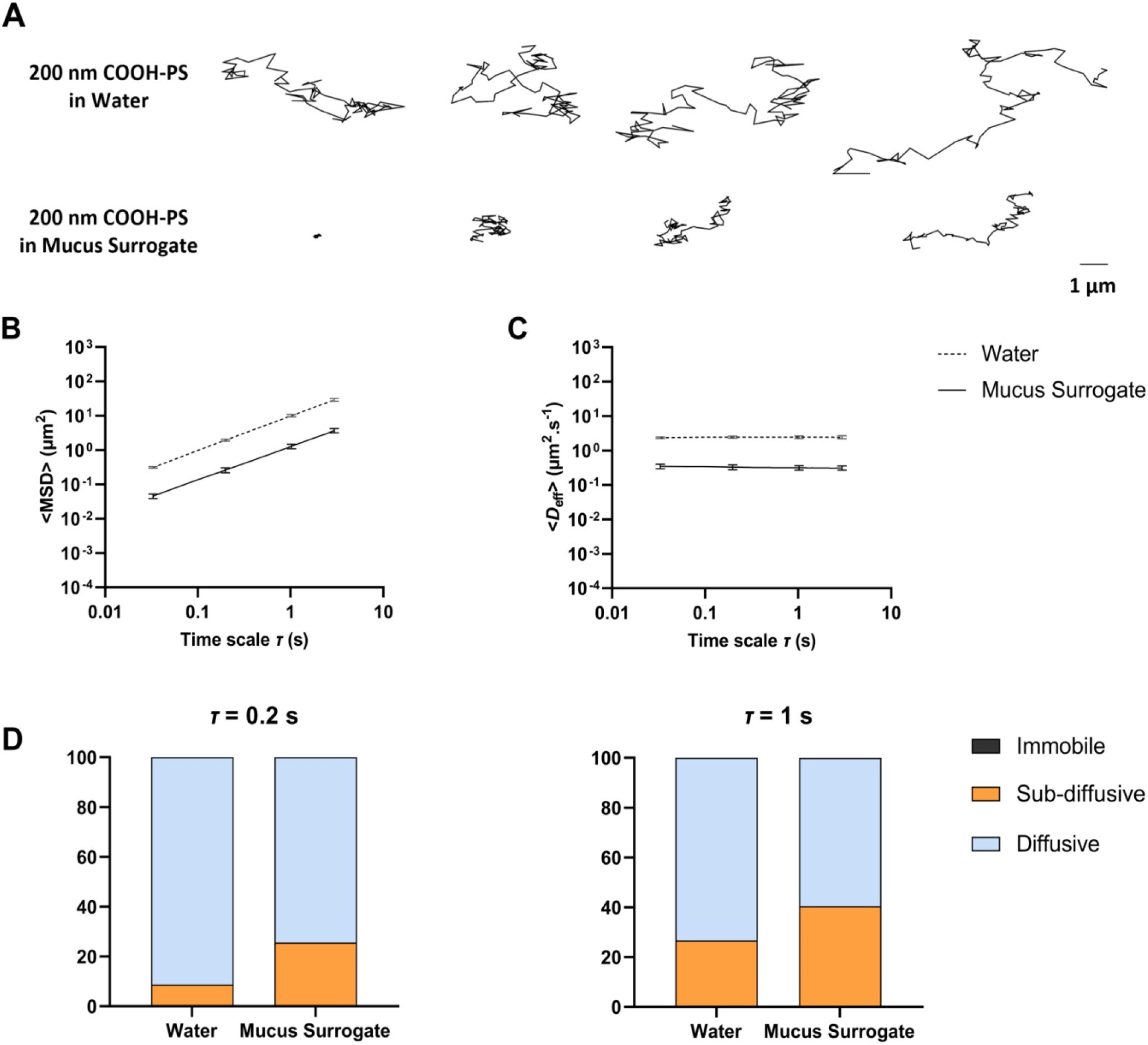
Transport behavior of 200 nm COOH-PS NPs in water or mucus surrogate containing 3% mucin. **(A)** Representative trajectories for a total duration of 3 sec. Values for **(B)** <MSD> and **(C)** <*D*_eff_> of 200 nm COOH-PS NPs in water or mucus surrogate containing 3% mucin as a function of time scale. Data presented as mean ± SD (*n* = 3). (D) Distribution of 200 nm COOH-PS NPs in water and mucus surrogate (3% mucin) according to transport mode for *τ* = 0.2 sec and *τ* = 1 sec (data presented as mean values; *n* = 3).

The values obtained for 200 nm COOH-PS NPs in mucus surrogate are in line with those reported by Crater and Carrier [34] using similar experimental conditions, but at a longer time scale value (10 sec). Nonetheless, transport hindrance of 200 nm COOH-PS NPs in either native cervicovaginal [5] or respiratory [41] mucus has been shown considerably higher (at least three orders of magnitude when compared to the theoretical mobility in water) than in the present work. Structural differences between native and surrogate media (obtained by simply reconstituting mucin in PBS) appear to largely justify such observations [34]. However, other possibilities may also help explaining higher mobility, namely the use of a short time scale that allowed faster particles to contribute with multiple trajectories (due to their easiness to offset the focus plane of the microscope during video capture) [42], or the high ionic strength of the mucus surrogate that can translate into diminished effective surface charge of NPs and, thus, reduce electrostatic interactions with mucin [43]. Average values for the anomalous diffusion exponent allowed classifying the ensemble transport of 200 nm COOH-PS NPs in water as the typical pure Brownian motion in viscous fluids (*α* = 0.98), while in the mucus surrogate as marginally sub-diffusive (*α* = 0.88) – likely due to the establishment of mild adhesive interactions with mucin and/or increased medium viscosity [44]. Nonetheless, this value is within the range of those described for similarly sized mucus-penetrating particles in native mucus [45].

Data from individual analysis also allowed classifying COOH-PS NPs in different classes of transport mode (**Figure 5D**). Virtually no particles were found to be immobile in either media. Nearly 90% of COOH-PS NPs were found diffusive in water (*τ* = 0.2 sec), contrasting with roughly 70% in mucus surrogate. Interestingly, the number of particles classified as diffusive decreased for *τ* = 1 sec, particularly in water. This has been previously justified by the increase in statistical uncertainty for higher values of lag time, since less contributions of MSDs are being considered for the calculation of its average [29]. The effect was more prominent in water since *τ* = 1 sec corresponds to more than 30% of the total length of the trajectories.

We further tested the applicability of MPTHub by changing experimental settings regarding the diameter (100 nm) and surface properties (PEG-modified using poloxamer 407 as coating agent) of fluorescent NPs, as well as medium composition (5% mucin). Summary data are presented in **Table 2**, while ensemble and individual distribution of MSD and *D*_eff_ are detailed in *Supplementary Information* (*Figures S12* to *S15*). Transport of 100 nm COOH-PS NPs in mucus surrogate containing 3% mucin was reduced as compared to their predictable behavior in water, but still within the diffusive range (*α* = 0.91). Hindrance was increased by around 2-fold when mucin content in the medium was increased from 3% to 5%, rendering marginally sub-diffusive behavior to particles. Non-covalent PEG-modification of 100 nm COOH-PS NPs impacted mobility in both mucus surrogates, featuring enhanced transport at both 3% and 5% mucin that is closer to the predicted behavior of similarly sized particles in water. These results are consistent with the established ability of densely PEG-modified NPs to present mucus-penetrating properties [46]. Still, the anomalous diffusion exponent at higher mucin concentration was nearly identical for modified and non-modified NPs at 5% mucin.

**Table 2.** Transport characterization of different 100 nm COOH-PS NPs in mucus surrogates containing either 3% or 5% mucin. Results are presented as mean values.

| Coating | Mucin content (w/w %) | $D_w/D_0$ | $\alpha$ |
| --- | --- | --- | --- |
| – | 3% | 6.0 | 0.91 |
| – | 5% | 12.1 | 0.86 |
| Poloxamer 407 | 3% | 1.7 | 0.98 |
| Poloxamer 407 | 5% | 4.9 | 0.88 |

## 4. Conclusions

MPTHub represents a user-friendly, free-to-use GUI allowing the quantitative study of particle transport in biorelevant media. The import of tracking data, information analysis, and export of relevant results is performed rapidly and robustly, allowing individual and ensemble transport characterization of fluorescent particles in mucus surrogates. Input data can be easily generated from simple video microscopy experiments and extracted using the popular ImageJ/Fiji software, while output data is complete and allows flexible visualization and edition according to the needs of specific users. Time savings achieved with the use of MPTHub, as compared to manual processing of tracking data, make it a powerful tool for high-throughput studies. However, additional work is required in order to allow further integration of the developed software with tracking algorithms (e.g. from ImageJ plugins) for direct trajectory extraction from videos, thus minimizing bias related with user input. Efforts for the incorporation of other features such as statistical processing and mathematical modeling are also underway and will be included in updated versions of MPTHub. Finally, future inclusion of the MPTHub in software performance comparison iterations [19] would be interesting for better understanding its advantages and limitations.

## Acknowledgments

Helena Almeida gratefully acknowledges Fundação para a Ciência e a Tecnologia, Portugal for financial support (2020.06264.BD fellowship). This work was financed by Portuguese funds through FCT - Fundação para a Ciência e a Tecnologia/Ministério da Ciência, Tecnologia e Ensino Superior in the framework of the project “Institute for Research and Innovation in Health Sciences” UID/BIM/04293/2019. The authors acknowledge the support of the following i3S Scientific Platforms: Biointerfaces and Nanotechnology, and Advanced Light Microscopy (member of the national infrastructure PPBI-Portuguese Platform of BioImaging and supported by POCI-01-0145-FEDER-022122).

