## Supplementary Information for "MPTHub: an open-source software for characterizing the transport of particles in biorelevant media"

**Table S1.** Details of the three computers used for testing the performance of MPTHub.

| Computer | Properties |
|---|---|
| #1 | Intel® Core™ i7-1165G7 processor (12M Cache, 2.80 GHz), 16 GB RAM |
| #2 | Intel® Core™ i7-4500U processor (4M Cache, 2.40 GHz), 16 GB RAM |
| #3 | Intel® Core™ i3-7020U processor (3M Cache, 2.30 GHz), 4 GB RAM |

| | A | B | C | D | E | F | G | |
|---|---|---|---|---|---|---|---|---|
| 1 | | | | | | | | |
| 2 | **Timescale (τ) (s)** | **MSD 1 (μm²)** | **MSD 2 (μm²)** | **MSD 3 (μm²)** | **MSD 4 (μm²)** | **MSD 5 (μm²)** | **MSD 6 (μm²)** | **MSD** |
| 3 | **0,033** | 0,3595162 | 0,335247965 | 0,358997906 | 0,321676024 | 0,333997556 | 0,346448755 | 0,324 |
| 4 | **0,066** | 0,748341197 | 0,711585943 | 0,805129822 | 0,663219512 | 0,739098928 | 0,696488509 | 0,700 |
| 5 | **0,099** | 1,120268097 | 1,174051432 | 1,304841505 | 0,921620463 | 1,121483404 | 1,091246289 | 1,082 |
| 6 | **0,132** | 1,469545014 | 1,622128414 | 1,852732019 | 1,169742825 | 1,502705168 | 1,509443649 | 1,439 |
| 7 | **0,165** | 1,766896682 | 2,115902296 | 2,444736938 | 1,445253865 | 1,923771135 | 1,929793572 | 1,81 |
| 8 | **0,198** | 2,051934254 | 2,575221646 | 3,10054382 | 1,741620877 | 2,342187519 | 2,372116057 | 2,149 |
| 9 | **0,231** | 2,386773102 | 3,061268762 | 3,822138155 | 2,060132619 | 2,830352796 | 2,805928233 | 2,534 |
| 10 | **0,264** | 2,658738134 | 3,526845485 | 4,650600635 | 2,367288709 | 3,340736842 | 3,214870477 | 2,95 |
| 11 | **0,297** | 2,958861547 | 4,01410966 | 5,510904781 | 2,69034476 | 3,831873943 | 3,625852242 | 3,291 |
| 12 | **0,33** | 3,247820116 | 4,489055587 | 6,452327422 | 3,045685978 | 4,386212756 | 4,001671855 | 3,587 |
| 13 | **0,363** | 3,581053401 | 4,951982046 | 7,454170951 | 3,409032259 | 4,904573731 | 4,363976118 | 3,847 |
| 14 | **0,396** | 3,899347909 | 5,388378578 | 8,51775556 | 3,806257712 | 5,429963842 | 4,739145238 | 4,128 |
| 15 | **0,429** | 4,227369462 | 5,815310817 | 9,589483376 | 4,188656076 | 6,008060872 | 5,100205105 | 4,429 |
| 16 | **0,462** | 4,566077196 | 6,272755398 | 10,6501328 | 4,588058194 | 6,624934629 | 5,481468795 | 4,763 |
| 17 | **0,495** | 4,919439938 | 6,703528742 | 11,68843967 | 5,00981054 | 7,215740261 | 5,873351624 | 5,082 |
| 18 | **0,528** | 5,267429448 | 7,163502192 | 12,78841263 | 5,461927387 | 7,903729865 | 6,283843286 | 5,382 |
| 19 | **0,561** | 5,570356636 | 7,629477234 | 13,94870602 | 5,815024209 | 8,63011697 | 6,643446367 | 5,676 |
| 20 | **0,594** | 5,80779689 | 8,134563816 | 15,08818778 | 6,206112829 | 9,399158759 | 7,045882493 | 6,035 |
| 21 | **0,627** | 6,055795755 | 8,678496593 | 16,24581773 | 6,547470114 | 10,17977257 | 7,4744327 | 6,390 |
| 22 | **0,66** | 6,37880983 | 9,26087019 | 17,42309556 | 6,822020101 | 10,99886098 | 7,937936049 | 6,738 |
| 23 | **0,693** | 6,660967315 | 9,788877614 | 18,52471306 | 7,147855432 | 11,8297804 | 8,422138196 | 7,098 |
| 24 | **0,726** | 7,017850546 | 10,38580921 | 19,74687562 | 7,476905524 | 12,66147328 | 8,934702678 | 7,512 |
| 25 | **0,759** | 7,345536493 | 11,03392893 | 21,00784292 | 7,78728221 | 13,47645427 | 9,418716461 | 7,894 |
| 26 | **0,792** | 7,624468151 | 11,71663269 | 22,24751252 | 8,064465729 | 14,3568269 | 9,888562771 | 8,259 |
| 27 | **0,825** | 7,903714939 | 12,41822614 | 23,43156142 | 8,277424859 | 15,32329787 | 10,30654166 | 8,601 |
| 28 | **0,858** | 8,197189283 | 13,10474539 | 24,54026547 | 8,413704862 | 16,35429489 | 10,72432633 | 8,929 |
| 29 | [illegible] | [illegible] | [illegible] | [illegible] | [illegible] | [illegible] | [illegible] | [illegible] |

Data | <MSD> vs Time | MSD vs Time | <Deff> vs Time | Deff vs Time

**Figure S1.** Example of generated 'Individual Particle Analysis' .xlsx file spreadsheet presenting raw analysis data after processing of input .csv files.

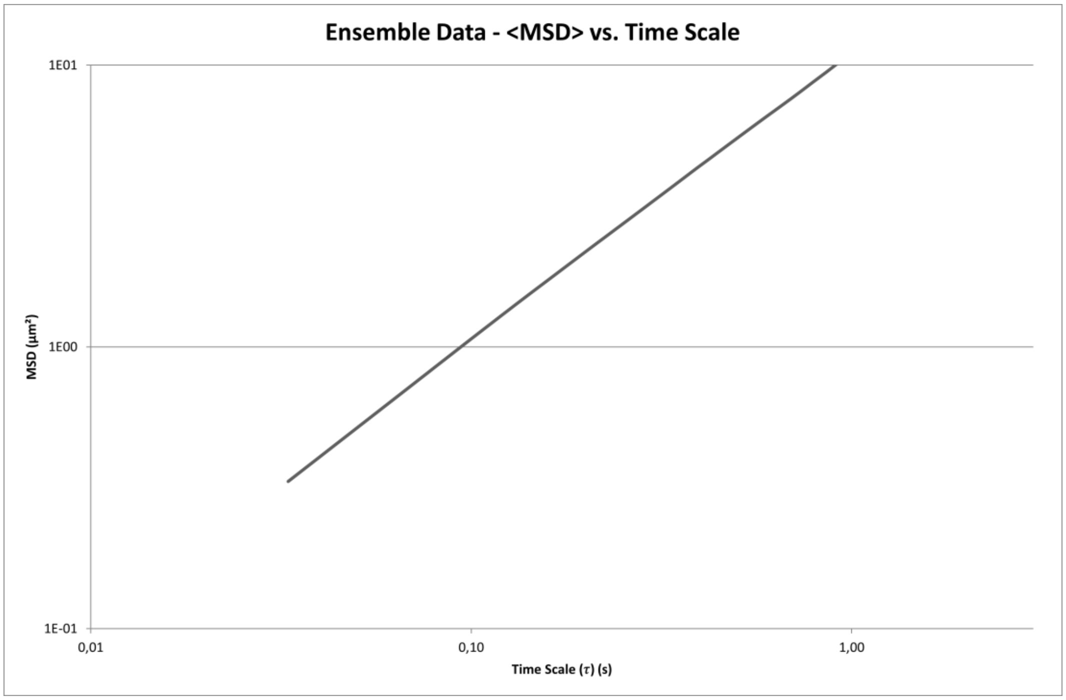


**Figure S2.** Example of a <MSD> vs. time scale graph generated in the 'Individual Particle Analysis' .xlsx file. Please note the logarithmic scales in both axes.

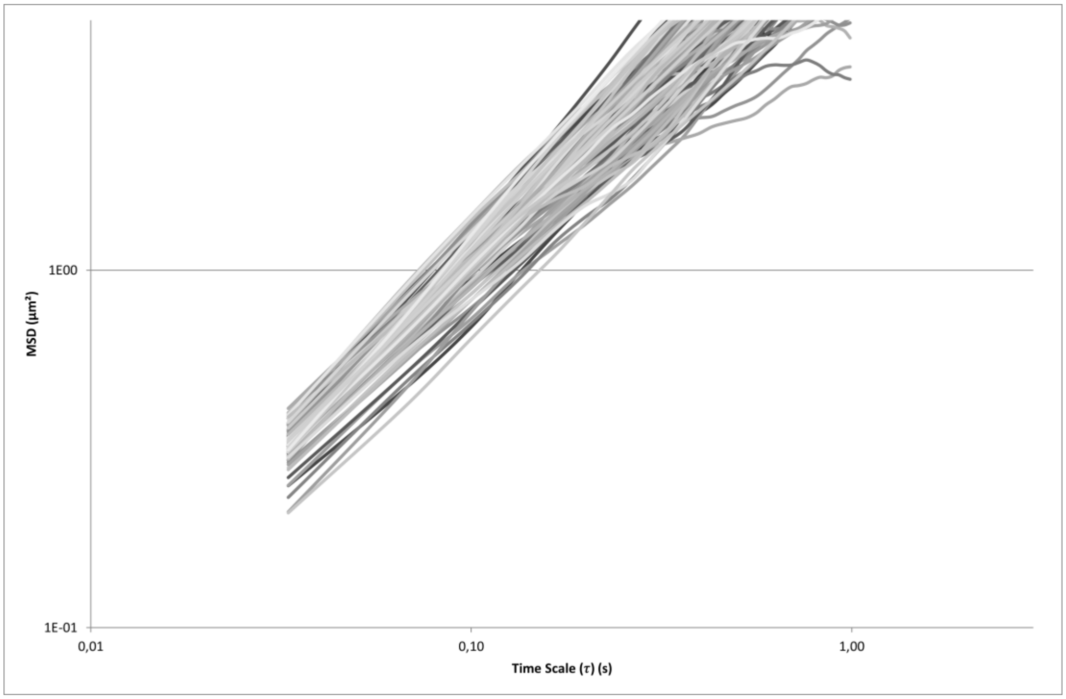


**Figure S3.** Example of a MSD vs. time scale graph generated in the 'Individual Particle Analysis' .xlsx file. Please note the logarithmic scales in both axes.

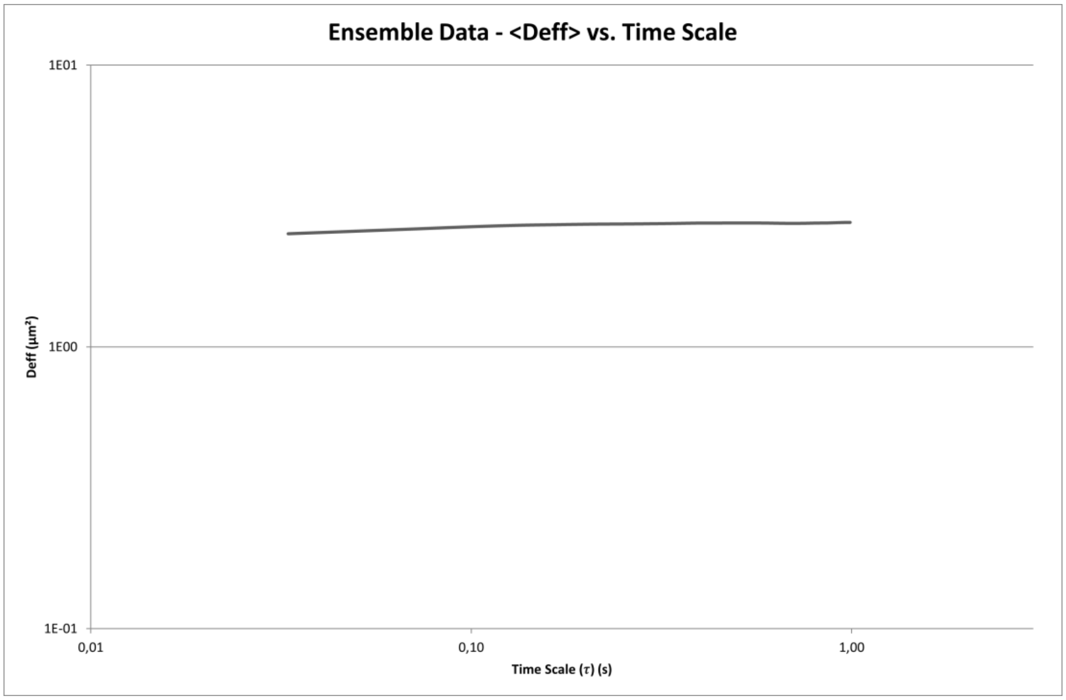


**Figure S4.** Example of a <$D_{eff}$> vs. time scale graph generated in the 'Individual Particle Analysis' .xlsx file. Please note the logarithmic scales in both axes.

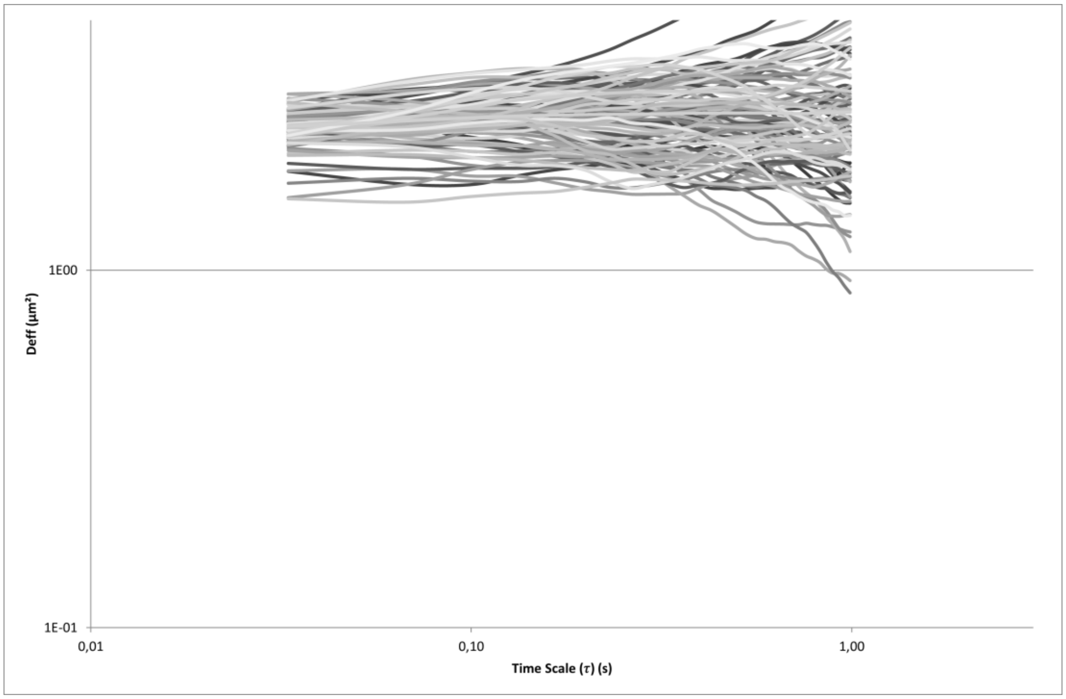


**Figure S5.** Example of a $D_{eff}$ vs. time scale graph generated in the 'Individual Particle Analysis' .xlsx file. Please note the logarithmic scales in both axes.

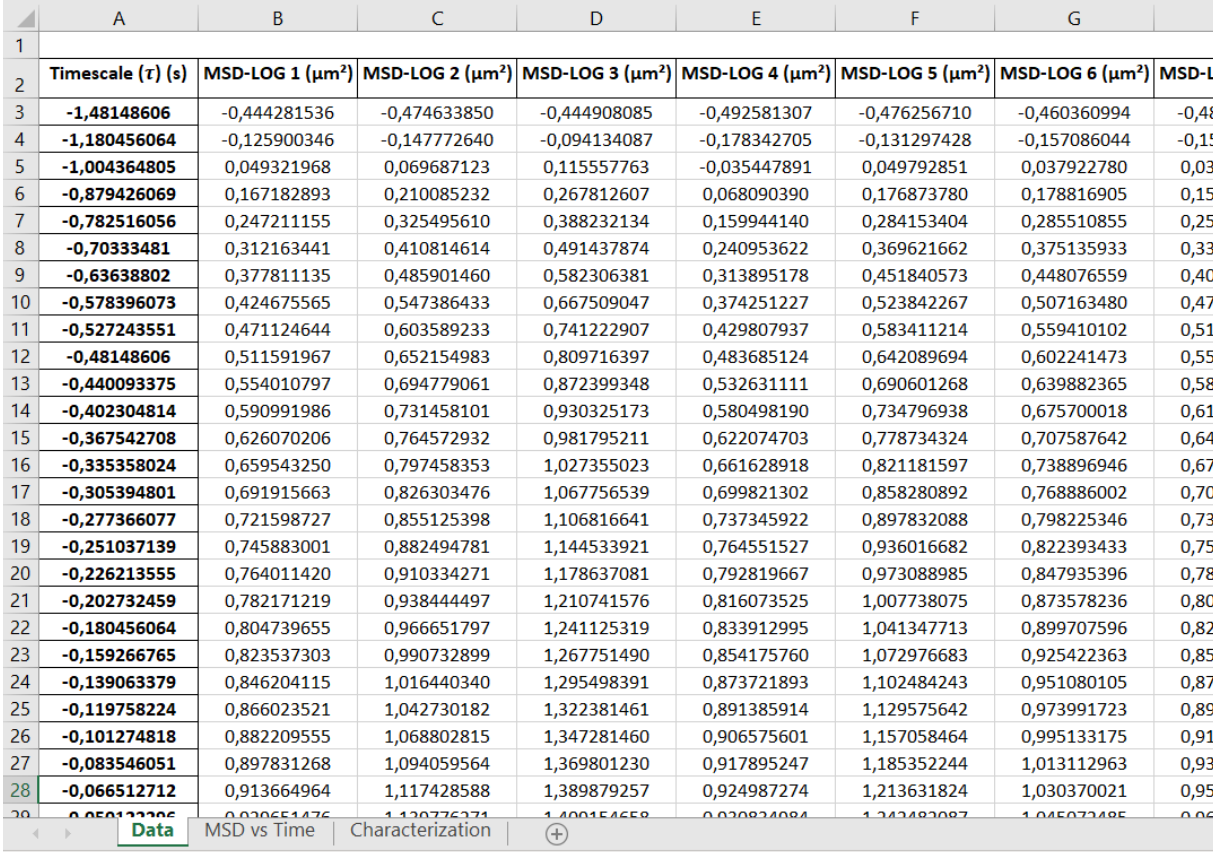

| | A | B | C | D | E | F | G | |
|---|---|---|---|---|---|---|---|---|
| 1 | | | | | | | | |
| 2 | **Timescale (τ) (s)** | **MSD-LOG 1 (μm²)** | **MSD-LOG 2 (μm²)** | **MSD-LOG 3 (μm²)** | **MSD-LOG 4 (μm²)** | **MSD-LOG 5 (μm²)** | **MSD-LOG 6 (μm²)** | **MSD-L** |
| 3 | **-1,48148606** | -0,444281536 | -0,474633850 | -0,444908085 | -0,492581307 | -0,476256710 | -0,460360994 | -0,48 |
| 4 | **-1,180456064** | -0,125900346 | -0,147772640 | -0,094134087 | -0,178342705 | -0,131297428 | -0,157086044 | -0,1 |
| 5 | **-1,004364805** | 0,049321968 | 0,069687123 | 0,115557763 | -0,035447891 | 0,049792851 | 0,037922780 | 0,03 |
| 6 | **-0,879426069** | 0,167182893 | 0,210085232 | 0,267812607 | 0,068090390 | 0,176873780 | 0,178816905 | 0,15 |
| 7 | **-0,782516056** | 0,247211155 | 0,325495610 | 0,388232134 | 0,159944140 | 0,284153404 | 0,285510855 | 0,25 |
| 8 | **-0,70333481** | 0,312163441 | 0,410814614 | 0,491437874 | 0,240953622 | 0,369621662 | 0,375135933 | 0,33 |
| 9 | **-0,63638802** | 0,377811135 | 0,485901460 | 0,582306381 | 0,313895178 | 0,451840573 | 0,448076559 | 0,40 |
| 10 | **-0,578396073** | 0,424675565 | 0,547386433 | 0,667509047 | 0,374251227 | 0,523842267 | 0,507163480 | 0,47 |
| 11 | **-0,527243551** | 0,471124644 | 0,603589233 | 0,741222907 | 0,429807937 | 0,583411214 | 0,559410102 | 0,51 |
| 12 | **-0,48148606** | 0,511591967 | 0,652154983 | 0,809716397 | 0,483685124 | 0,642089694 | 0,602241473 | 0,55 |
| 13 | **-0,440093375** | 0,554010797 | 0,694779061 | 0,872399348 | 0,532631111 | 0,690601268 | 0,639882365 | 0,58 |
| 14 | **-0,402304814** | 0,590991986 | 0,731458101 | 0,930325173 | 0,580498190 | 0,734796938 | 0,675700018 | 0,61 |
| 15 | **-0,367542708** | 0,626070206 | 0,764572932 | 0,981795211 | 0,622074703 | 0,778734324 | 0,707587642 | 0,64 |
| 16 | **-0,335358024** | 0,659543250 | 0,797458353 | 1,027355023 | 0,661628918 | 0,821181597 | 0,738896946 | 0,67 |
| 17 | **-0,305394801** | 0,691915663 | 0,826303476 | 1,067756539 | 0,699821302 | 0,858280892 | 0,768886002 | 0,70 |
| 18 | **-0,277366077** | 0,721598727 | 0,855125398 | 1,106816641 | 0,737345922 | 0,897832088 | 0,798225346 | 0,73 |
| 19 | **-0,251037139** | 0,745883001 | 0,882494781 | 1,144533921 | 0,764551527 | 0,936016682 | 0,822393433 | 0,75 |
| 20 | **-0,226213555** | 0,764011420 | 0,910334271 | 1,178637081 | 0,792819667 | 0,973088985 | 0,847935396 | 0,78 |
| 21 | **-0,202732459** | 0,782171219 | 0,938444497 | 1,210741576 | 0,816073525 | 1,007738075 | 0,873578236 | 0,80 |
| 22 | **-0,180456064** | 0,804739655 | 0,966651797 | 1,241125319 | 0,833912995 | 1,041347713 | 0,899707596 | 0,82 |
| 23 | **-0,159266765** | 0,823537303 | 0,990732899 | 1,267751490 | 0,854175760 | 1,072976683 | 0,925422363 | 0,85 |
| 24 | **-0,139063379** | 0,846204115 | 1,016440340 | 1,295498391 | 0,873721893 | 1,102484243 | 0,951080105 | 0,87 |
| 25 | **-0,119758224** | 0,866023521 | 1,042730182 | 1,322381461 | 0,891385914 | 1,129575642 | 0,973991723 | 0,89 |
| 26 | **-0,101274818** | 0,882209555 | 1,068802815 | 1,347281460 | 0,906575601 | 1,157058464 | 0,995133175 | 0,91 |
| 27 | **-0,083546051** | 0,897831268 | 1,094059564 | 1,369801230 | 0,917895247 | 1,185352244 | 1,013112963 | 0,93 |
| 28 | **-0,066512712** | 0,913664964 | 1,117428588 | 1,389879257 | 0,924987274 | 1,213631824 | 1,030370021 | 0,95 |
| 29 | [illegible] | [illegible] | [illegible] | [illegible] | [illegible] | [illegible] | [illegible] | [illegible] |



**Figure S6.** Example of generated 'Transport Mode Characterization' .xlsx file spreadsheet presenting raw analysis data after processing of input .csv files.

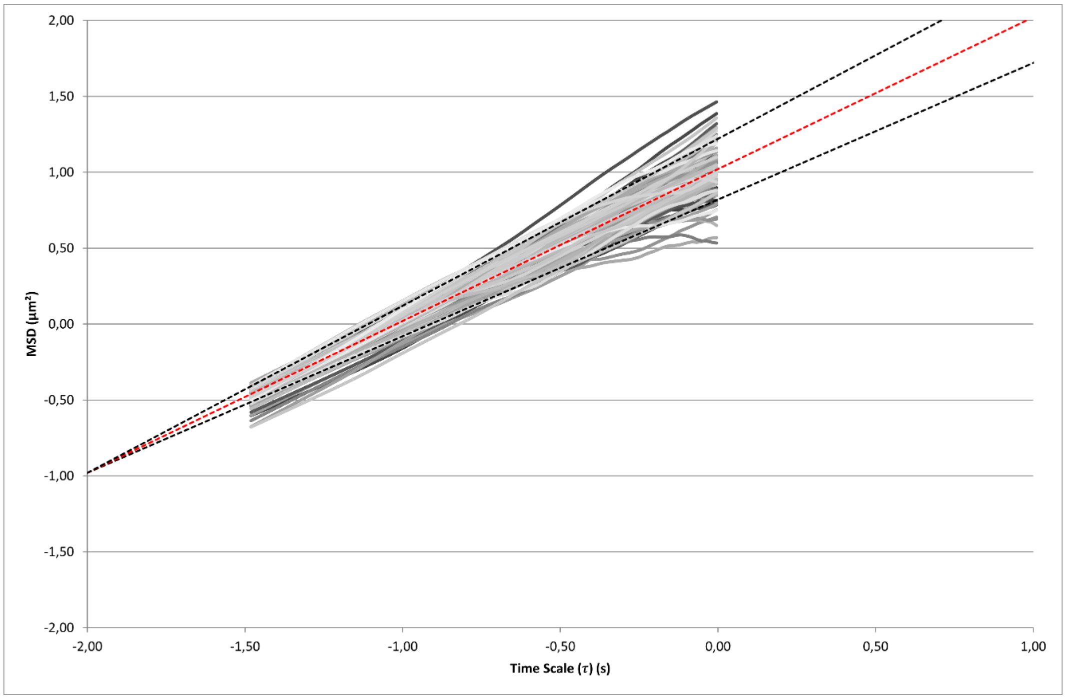


**Figure S7.** Example of a MSD vs. time scale graph generated in the 'Transport Mode Characterization' .xlsx file. Dashed lines presenting different slopes (0.9, 1.0 and 1.1) are included for guidance. Please note the linear scales in both axes.

| | A | B | C | D | E | F |
|---|---|---|---|---|---|---|
| 1 | **Slopes** | **Slopes (Excel)** | **$R^2$ (Excel)** | **Transport Mode** | **Slope** | **Count** |
| 2 | 0,946003222 | 0,946003222 | 0,999286258 | Immobile | 0,0-0,199 | 0 |
| 3 | 1,113317046 | 1,113317046 | 0,999057535 | Sub-diffusive | 0,2-0,899 | 19 |
| 4 | 1,340696417 | 1,340696417 | 0,998223359 | Diffusive | 0,9-1,199 | 76 |
| 5 | 1,028013443 | 1,028013442 | 0,99645975 | Active | 1,2+ | 7 |
| 6 | 1,225965352 | 1,225965352 | 0,994306885 | | | |
| 7 | 1,044505004 | 1,044505004 | 0,999415628 | | | |
| 8 | 0,990424520 | 0,99042452 | 0,998715094 | <slope> = | 0,997907128 | |
| 9 | 0,936409573 | 0,936409573 | 0,996765191 | N = | 102 | |
| 10 | 0,926102028 | 0,926102028 | 0,992990995 | STD = | 0,134222422 | |
| 11 | 0,820440377 | 0,820440377 | 0,979635679 | | | |
| 12 | 1,014209825 | 1,014209824 | 0,993315373 | | | |
| 13 | 0,802135139 | 0,802135139 | 0,989021957 | | | |
| 14 | 0,978573239 | 0,978573239 | 0,997081521 | | | |
| 15 | 1,040239918 | 1,040239918 | 0,992301314 | | | |
| 16 | 1,083384612 | 1,083384612 | 0,998257775 | | | |
| 17 | 0,946599831 | 0,946599831 | 0,996841926 | | | |
| 18 | 1,132902333 | 1,132902333 | 0,997483036 | | | |
| 19 | 1,061521906 | 1,061521906 | 0,991163249 | | | |
| 20 | 0,880342849 | 0,88034285 | 0,929800207 | | | |
| 21 | 1,255023774 | 1,255023774 | 0,99123038 | | | |
| [illegible] | [illegible] | [illegible] | [illegible] | | | |

**Figure S8**. Example of a spreadsheet containing individual and ensemble data summary in the 'Transport Mode Characterization' .xlsx file.

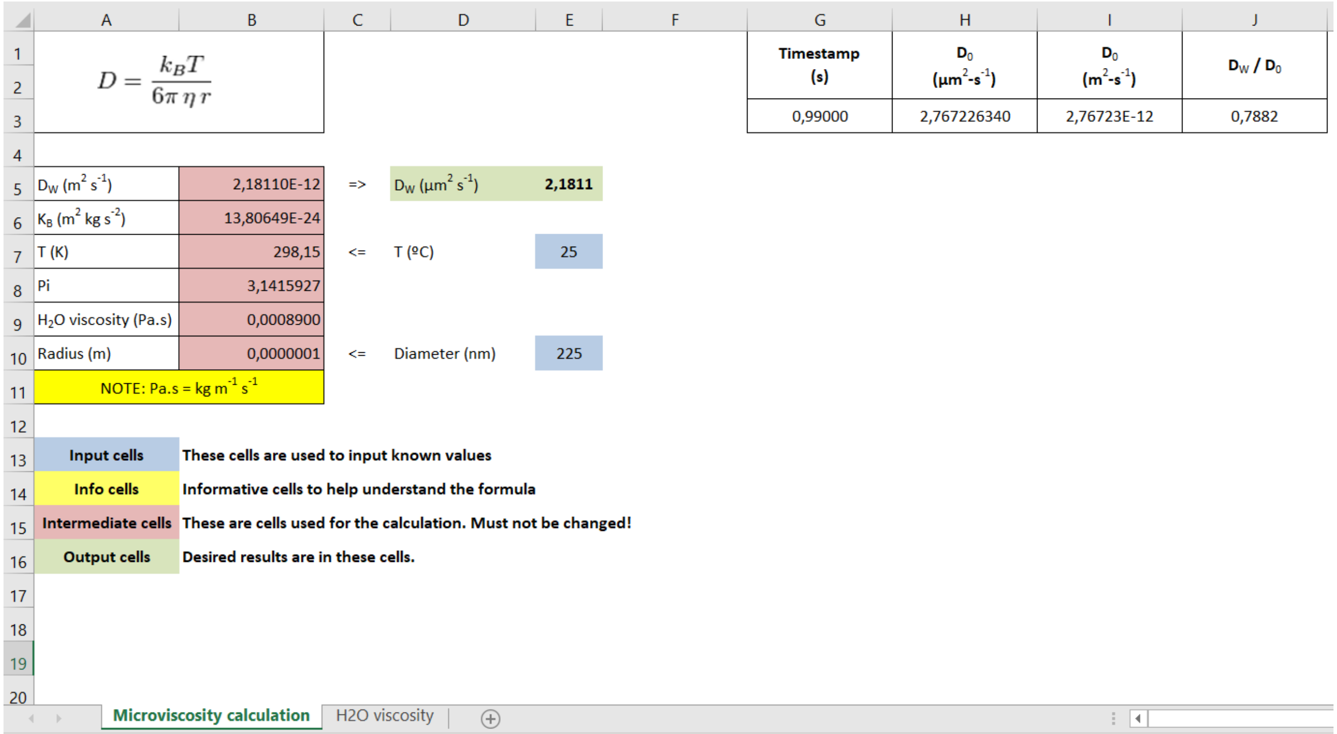


**Figure S9.** Example of generated 'Stokes-Einstein Calculations' .xlsx file spreadsheet generated after processing of input .csv files.

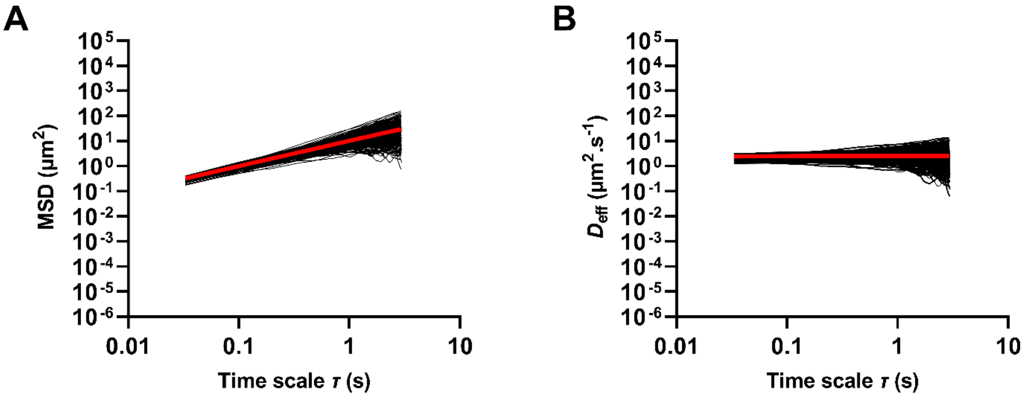


**Figure S10.** Individual **(A)** MSD and **(B)** $D_{eff}$ of 200 nm COOH-PS NPs (≥ 100 tracked particles per experiment; *n* = 3) in water as a function of time scale. Ensemble values are also presented in red in both graphs.

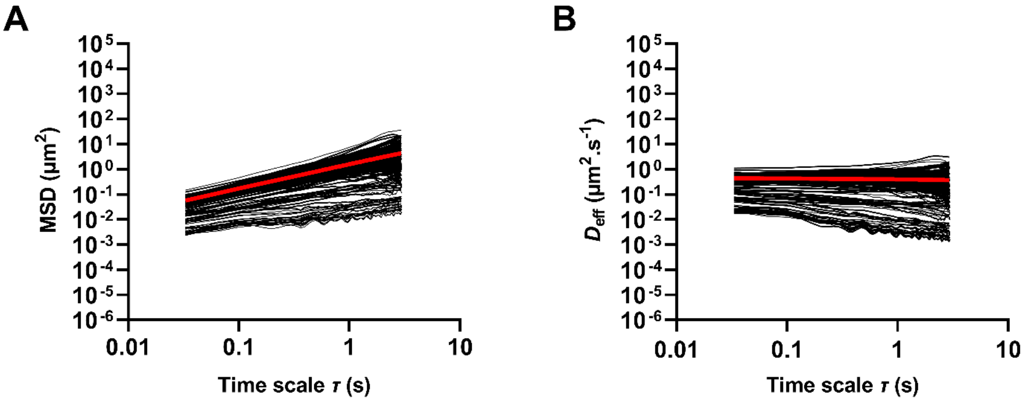


**Figure S11.** Individual **(A)** MSD and **(B)** $D_{eff}$ of 200 nm COOH-PS NPs (≥ 100 tracked particles per experiment; *n* = 3) in mucus surrogate containing 3% mucin as a function of time scale. Ensemble values are also presented in red in both graphs.

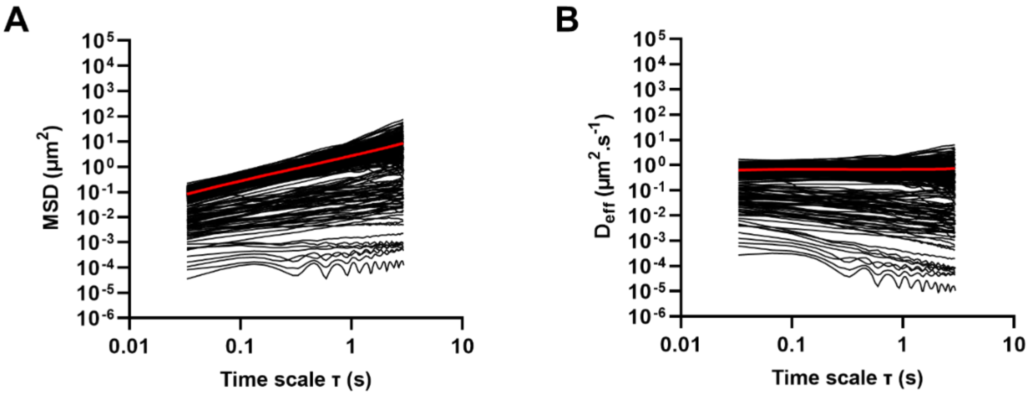


**Figure S12.** Individual **(A)** MSD and **(B)** $D_{eff}$ of 100 nm COOH-PS NPs (≥ 100 tracked particles per experiment; *n* = 3) in mucus surrogate containing 3% mucin as a function of time scale. Ensemble values are also presented in red in both graphs.

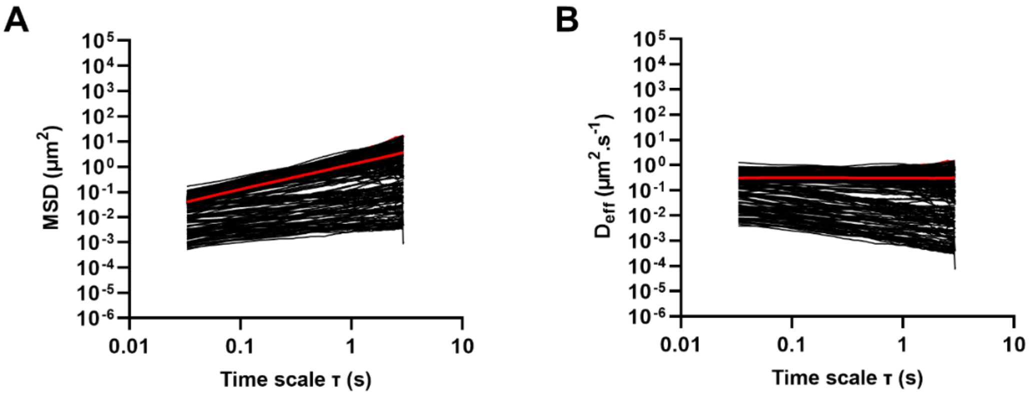


**Figure S13.** Individual **(A)** MSD and **(B)** $D_{eff}$ of 100 nm COOH-PS NPs (≥ 100 tracked particles per experiment; $n$ = 3) in mucus surrogate containing 5% mucin as a function of time scale. Ensemble values are also presented in red in both graphs.

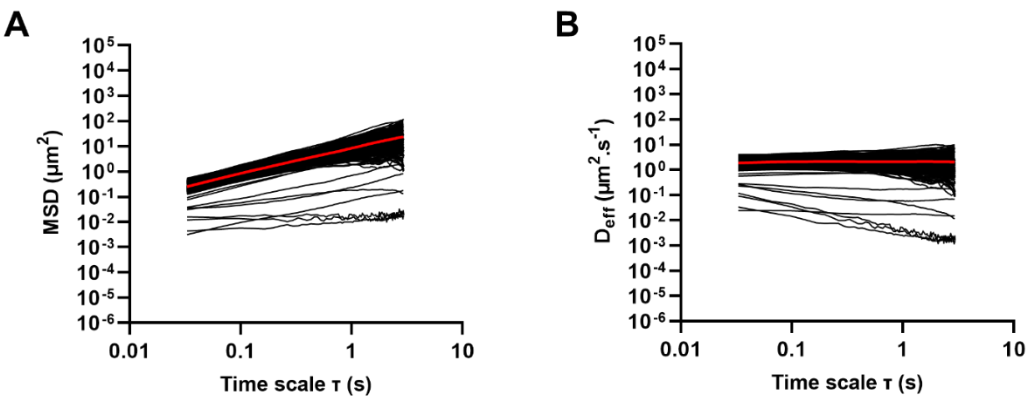


**Figure S14.** Individual **(A)** MSD and **(B)** $D_{eff}$ of 100 nm PEG-modified COOH-PS NPs (≥ 100 tracked particles per experiment; *n* = 3) in mucus surrogate containing 3% mucin as a function of time scale. Ensemble values are also presented in red in both graphs.

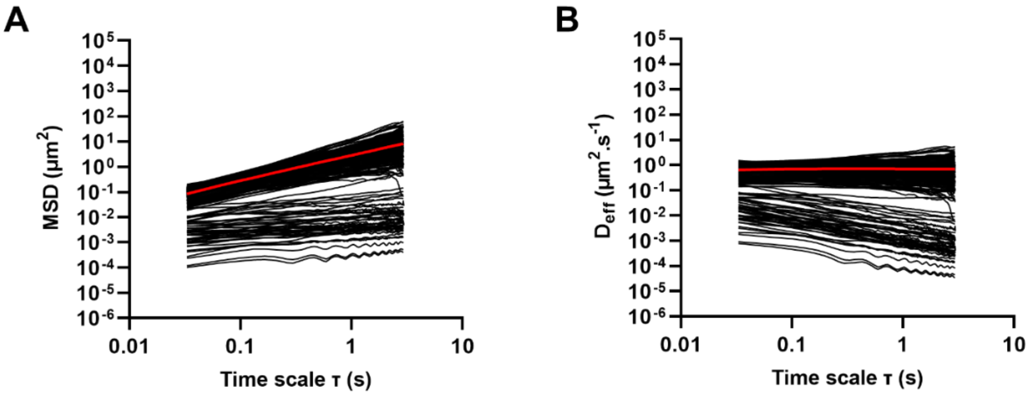


**Figure S15.** Individual **(A)** MSD and **(B)** $D_{eff}$ of 100 nm PEG-modified COOH-PS NPs (≥ 100 tracked particles per experiment; *n* = 3) in mucus surrogate containing 5% mucin as a function of time scale. Ensemble values are also presented in red in both graphs.
